# Age and sex modulate the variability of neural responses to naturalistic videos

**DOI:** 10.1101/089060

**Authors:** Agustin Petroni, Samantha Cohen, Lei Ai, Nicolas Langer, Simon Henin, Tamara Vanderwal, Michael P. Milham, Lucas C. Parra

## Abstract

Neural development is generally marked by an increase in the efficiency and diversity of neural processes. In a large sample (N = 114) of children and adults with ages ranging from 5 −44 years, we investigated the neural responses to naturalistic video stimuli. Videos from both real-life classroom settings and Hollywood feature films were used to probe different aspects of attention and engagement. For all stimuli, older ages were marked by more variable neural responses. Variability was assessed by the inter-subject correlation of evoked electroencephalographic (EEG) responses. Young males also had more variable responses than young females. These results were replicated in an independent cohort (N = 303). When interpreted in the context of neural maturation, we conclude that neural function becomes more variable with maturity, at least in during the passive viewing of real-world stimuli.

**Significance Statement:** Naturalistic videos were used to measure how a large sample of children and adults process environmentally meaningful stimuli. As age increased, neural responses were more variable, and females responded more variably than males - a difference that disappeared with age. These results are consistent with developmental theories positing that neural variability increases with maturation, and that neural maturation typically occurs earlier in females. This is the first study to investigate neural variability under naturalistic conditions in a developmental sample.

## Introduction

Over the course of development, the accuracy and stability of behaviors generally increase. This performance improvement is typically accompanied by a seemingly paradoxical increase in the variability of neural responses (Grady, 2012; Dinstein et al., 2015). More variable electroencephalographic (EEG) responses among adults are associated with lower reaction time variability and higher recognition accuracy (Mcintosh et al., 2008). Neural variability often presents as an increase in the complexity of neural responses. This may be due in part to a developmental increase in the repertoire of possible brain states (Vakorin et al., 2013) and this increase in complexity may underlie the integration between distributed neural populations (Vakorin et al., 2011). EEG signal complexity becomes elevated in late adolescence and is also elevated in females relative to males at this stage, indicating that females may reach neural maturity prior to males (Anokhin et al., 2000). Anatomical studies generally support the notion that females reach neural maturity prior to males (Giedd et al., 1999; Lenroot and Giedd, 2006; Lenroot et al., 2007; Marsh et al., 2008).

Neural variability does not always accompany proficient behavior, however. Both theta band coherence, a performance monitoring measure, and behavior are more variable in children (Papenberg et al., 2013). This suggests that neural variability does not always increase with maturation. For adults, the variability in “functional connectivity” between different networks measured with fMRI is elevated during rest and decreases during a cognitive task. The reverse is true for children, whose brains become more variable during the task, and their performance is expectedly lower than adults (Hutchison and Morton, 2015).

Recently, responses to naturalistic narrative stimuli have been used to examine how variability in behavior and neural activity change with development. Using *Sesame Street* videos, it has been found that infants have more variable eye gaze patterns than adults (Kirkorian et al., 2012), and adults have more broadly similar neural responses than children (Cantlon and Li, 2013). While adults respond to *Sesame Street* more similarly in many parietal and frontal regions, children respond more similarly in a specific region in the superior temporal cortex (Cantlon and Li, 2013). Generally, from ages 18-88, as humans age, responses to videos increase in variability (Campbell et al., 2015). Taken together, these studies demonstrate that neural variability changes with age. The trajectory of this change depends on multiple factors including the metric of neural variability, the developmental stage sampled, and the brain region(s) of interest.

Here, EEG was recorded from subjects with ages ranging from 5 - 44 years as they were presented with both naturalistic (Dmochowski et al., 2012, 2014) and conventional stimuli. To assess neural variability, the level of similarity across subjects was assessed with the inter-subject correlation (ISC) of responses evoked by the stimuli. ISC of the EEG is indicative of attention, engagement, and memory in healthy adults (Dmochowski et al., 2014; Cohen and Parra, 2016; Ki et al., 2016; Cohen et al., 2017). We found that neural responses, indexed by ISC, become more variable with age. Among children, females have more variable neural responses than males. This increase in variability is not due to a decrease in evoked response magnitude, and was reproduced in two independent cohorts consisting of 114 and 303 individuals. These results are consistent with theories positing that development coincides with an increased repertoire of neural representations (Mcintosh et al., 2008), and the sex differences are consistent with the idea that young males are less neurally mature than young females (Giedd et al., 1999; Lenroot and Giedd, 2006; Lenroot et al., 2007; Marsh et al., 2008). Importantly, this is the first EEG study to report a measure of across-subject neural similarity with clear age and sex effects.

## Methods

### Subjects

In the main study ages ranged from 6 to 44 years old (N = 114, 14.2 +/- 8.0 years old, 46 females, see Figure 1A for a full age and sex distribution) as part of the Child Mind Institute - Multimodal Resource for Studying Information Processing in the Developing Brain (MIPDB; http://fcon_1000.projects.nitrc.org/indi/cmi_eeg/; (Langer et al., 2017)). In the replication study ages ranged from 5 to 21 years old (N = 303, 11.3 +/- 3.9 years old, 135 females, see figure 1B for a full age and sex distribution). This data was obtained from the Child Mind Institute Healthy Brain Network (CMI-HBN; http://fcon_1000.projects.nitrc.org/indi/cmi_healthy_brain_network/; (Alexander et al., 2017)). All experiments were performed in accordance with relevant guidelines and regulations. The study was reviewed and approved by the Chesapeake Institutional Review Board. All subjects presented with normal or corrected to normal vision.

**Figure 1:**
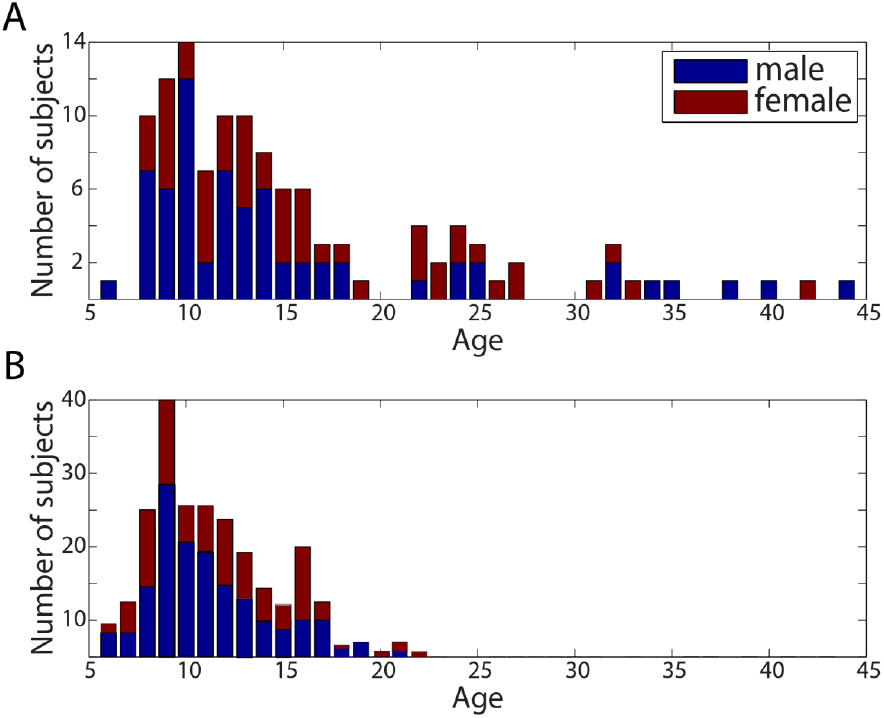
Age and sex distributions for the main study (A) and replication study (B). A: Subjects in the main study (N=114) had data for all of the stimuli. B: Subjects in the replication study (N=303) had only three stimuli (Wimpy, Fract, DesMe) and contribute to the results in Figure 9.

### Stimuli

Engaging, naturalistic videos were the primary stimuli. Specific videos were selected because they contained content relevant to social cognition, classroom anxiety, and attention. The videos ranged in engagement level. The weakly engaging videos contained educational content or depicted learning scenarios: *Fun with Fractals* (Fract, MIT), a cartoon that explains fractals with examples (4m 34s), *How to improve at Simple Arithmetic* (Arith, E-How), in which a math teacher in a typical educational setting explains addition and multiplication (1m 30s), and *Pre-Algebra Class* (StudT, Pearson Education)), showing an interaction between two students and a teacher (StudT, for student-teacher interaction) during math problem solving (1m 40s). The most engaging videos were from conventional cinema: *Diary of a Wimpy Kid* (Wimpy, Universal Pictures), a movie about a preteen starting middle school (1m 57s), and *Despicable Me* (DesMe, Universal Pictures), which contains infant and toddler characters and emphasizes social interactions (2m 51s). URLs for all of the videos and images for all of the stimuli are in the legend of Figure 2. While the main cohort contains data from all stimuli, the replication cohort only had three stimuli: Wimpy, Fract, and DesMe. The variability of the neural responses to these videos was measured across subjects using the inter-subject correlation (ISC) of evoked responses (see below). As a control condition, a “Rest” condition, during which subjects sat with their eyes-closed for 4m 20s, was also analyzed. This period establishes the baseline level of ISC, as no time-aligned stimulus entraining neural activity across subjects was presented. Another condition with a low engagement level was “Flash.” During this stimulus a black and white grating pattern that flashed at 25 Hz was presented for three minutes, thus synchronously stimulating neural activity across subjects (see Steady State Visual Evoked Potentials (SSVEP) Methods section and Figure 2). This stimulus elicits steady state evoked potentials (Vanegas et al., 2015) and was included to explore the extent to which ISC is driven by low level evoked responses.

**Figure 2:**
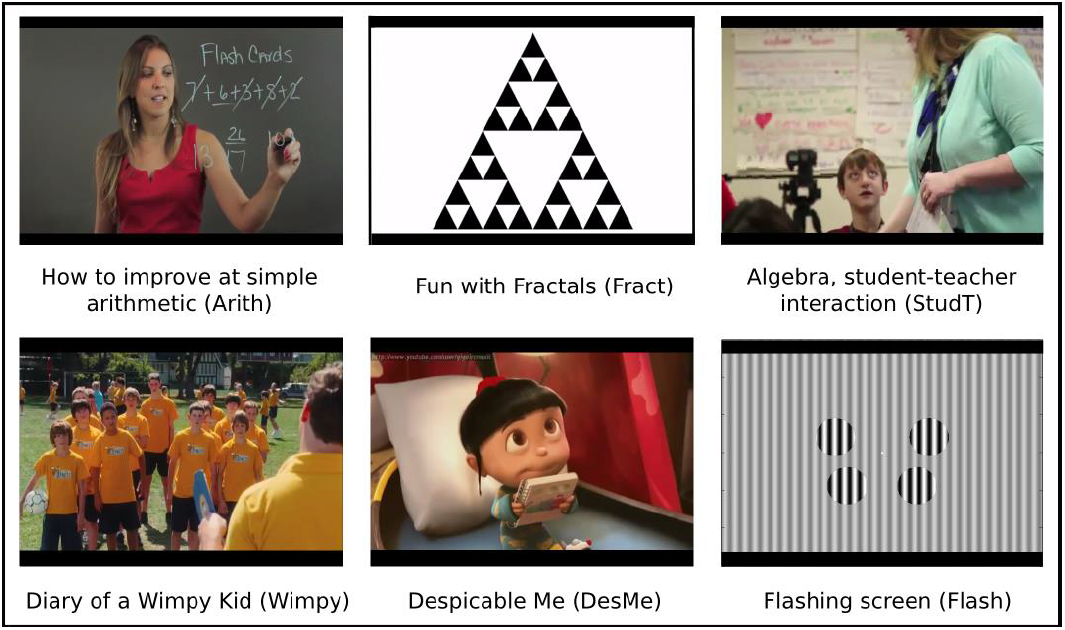
Still images from all stimuli except Rest which contained no stimulus. Video URLs, and total duration (time segment used is also included if applicable): Despicable Me (DesMe): http://www.youtube.com/watch?v=HNXxJIhVALI (2:51), The Diary of a Wimpy Kid Trailer (Wimpy): http://www.youtube.com/watch?v=7ZVEIgPeDCE (1:57), Fun with Fractals (Fract): http://www.youtube.com/watch?v=XwWyTts06tU (4:40), How to Improve at Simple Arithmetic (Arith): http://www.youtube.com/watch?v=pHoE7AMtXcA (1:40), Pre-Algebra Class (StudT): https://www.youtube.com/watch?v=wyClt5S0gF8 (segment: 10:59 - 12:27, total duration: 1:28)

### Procedure

While seated in a dimly lit room wearing an EEG net, subjects watched a series of short videos in a pseudorandom order. Stimuli were presented on a 17-inch CRT monitor (SONY Trinitron Multiscan G220, display dimensions 330×240 mm, resolution 800×600 pixels, vertical refresh rate of 100 Hz). Note that some subjects did not experience all stimuli due to time limitations (Langer et al., 2017). Additionally, as explained below, poor data quality for some recordings caused additional data loss. For the replication study, only three condition were used: Wimpy, Fract and DesMe.

### EEG recordings and preprocessing

EEG recordings were performed with an EGI Clinical Geodesic 128 channel system (Electrical Geodesic Inc, Eugene, OR). Of the 128 channels recorded, 105 constituted the EEG recording and 11 represented EOG channels used for eye movement artifact removal. The remaining channels, mainly recording from the neck and face, were discarded. First, noisy channels were selected by visual inspection and replaced with by zero valued samples, thus eliminating those channels’ contribution in subsequent calculations of covariance matrices. Recordings, initially at 500 Hz, were then downsampled to 125 Hz, high-pass filtered at 1 Hz, and notch filtered between 59 and 61 Hz with a 4th-order Butterworth filter. Eye artifacts were removed by linearly regressing the EOG channels from the scalp EEG channels (Parra et al., 2005). Next, a robust Principal Components Analysis (PCA) algorithm, the inexact Augmented Lagrange Multipliers Method (Lin et al., 2013), was used to removed sparse outliers from the data following Ki et al (2016). Briefly, robust PCA recovers a low-rank matrix, A, from a corrupted data matrix D = A + E, where some entries of the additive errors E may be arbitrarily large. Finally, individual recordings for some stimuli were discarded on the basis of visual inspection because they remained noisy after both automatic and manual noise removal. This was necessary because these subjects exhibited profound movement artifacts and/or the saline used for the recordings dried out. All signal processing was performed offline using MATLAB software (MathWorks, Natick, MA, USA).

### Inter-Subject Correlation (ISC)

As variability is the inverse of similarity, we measured the similarity of evoked EEG responses across subjects. This approach has been used extensively to study concerted, inter-subject changes in blood-oxygen level dependent (BOLD) signal in fMRI (Hasson et al., 2004, 2009; Kauppi et al., 2010), and has been adapted to leverage the improved time resolution facilitated by EEG. To determine the neural similarity across subjects responding to the same stimulus (or in the same condition, in the case of Rest) the inter-subject correlation (ISC) of the EEG signal was computed, as described previously (Dmochowski et al., 2012, 2014; Cohen and Parra, 2016; Ki et al., 2016). ISC assesses the level of correlation in the EEG among a group of subjects as they respond to the same stimulus. Larger ISC values imply more similarity in fast EEG responses across subjects (< 1s). This indicates that the signals are more reliable due to decreased inter-subject variability. It has also been found that subjects who pay more attention to the stimulus have higher ISC values (Ki et al., 2016). An advantage of the technique is that the stimulus need only be presented once to each subject because evoked responses are compared across individuals. As repeated trials are unnecessary, responses are more similar to natural situations in which people experience uniquely presented novel stimuli. Additionally, in contrast to event related potentials, the technique can be applied to continuous and dynamic natural stimuli without the need for specific event markers (Ben-Yakov et al., 2012). As such, the approach is data-driven (i.e., *a priori)* both spatially and temporally.

ISC utilizes correlated component analysis to identify linear combinations of EEG electrodes that capture most of the correlation across subjects (Dmochowski et al., 2012). Correlated component analysis is similar to principal component analysis (PCA) except that rather than maximizing variance within one dataset, it selects projections that maximize the correlation between multiple datasets. These projections can be thought of as virtual sensors (or component sources) of activity that are optimized to capture most of the correlation between subjects. Following previous research, we use the three components that represent most of the correlation across subjects. These components, can be optimized for all subjects together, or for a subset of the entire cohort. The subsets used in this paper are stimulus (Wimpy, DesMe, Fract, Arith, StudT, Flash, and Rest), age group (young vs old), sex (male vs female), and sex and age group combined (young-male, young-female, old-male, and old-female). ISC components are computed within subsets of the entire sample to allow for potential differences in the spatial distribution of activity across groups, although the spatial patterns are largely consistent (Figure 8).

To calculate the ISC for individual subjects as they respond to the same condition as their peers, the correlation between each individual’s EEG responses and the responses from all other individuals is calculated (Cohen and Parra, 2016; Ki et al., 2016). The ISC values reported throughout the paper are this measure of how well each individual correlates with the others. The components used to compute this subject-specific ISC value are either computed across all subjects or within the subgroups listed above (divided by either stimulus, age, sex, or age and sex). A fuller description of the mathematical basis of the analysis is available in Ki et al. (2016) and Cohen and Parra (2016). A simplified template for the code to compute the correlated components and the ISC for individual subjects is available at http://www.parralab.org/isc/

### Steady state visual evoked potentials (SSVEPs)

To determine the strength of low-level sensory evoked responses across individuals, we leveraged the steady state visual evoked potential (SSVEP) paradigm (Flash) that was part of the data collection effort (Langer et al., 2017). Stimulus and analysis followed established techniques (Vanegas et al., 2015). Briefly, the stimulus consisted of four circular ‘foreground’ stimuli (vertical grating, radius 2°) that were flickered on-and-off at 25 Hz and embedded in a static ‘background’ grating, which is known to generate reliable SSVEPs (Vanegas et al., 2015). This stimulus was presented in trials of 2.4 s duration with inter-trial intervals of 1s which included a fixation cross presented for 0.5 s. The stimuli were presented in several conditions that varied in their contrast and in the phase relationship between the foreground and the background. A total of 128 trials were present (12 conditions total: four foreground contrasts - 0% 30%, 60% and 100%, and three background conditions - parallel phase, orthogonal phase, and no surround stimuli). Artifacts were rejected by removing trials for which the power in any electrode exceeded more than three standard deviations above the mean. EOG activity was regressed out of the EEG, as described above. The initial 200ms of each trial was removed to eliminate the onset of the visual evoked response. Data were Fourier transformed for each trial, power in the 25 Hz band was extracted, and then averaged across all trials, regardless of condition (thus ignoring details of the foreground-background interaction). Since the EEG activity measured with this paradigm is known to be dominated by primary visual cortex (V1) responses, power was averaged over the five most relevant occipital electrodes (O1-O5; (Vanegas et al., 2015)).

## Results

We sought to determine whether and how the variability of EEG differs across age and gender in children and adults ranging from 6 – 44 years of age. To assess the variability in EEG signals across subjects, the intersubject correlation (ISC) between individuals and their peers was assessed in response to both naturalistic videos and artificial stimuli. ISC can be thought of as a measure of the similarity of neural responses (Dmochowski et al., 2012). If subjects respond more similarly to their peers, they will have a larger ISC value, which indicates that they have a less variable neural response.

### Intersubject correlation varies between stimuli

The stimuli used ranged in engagement level from no stimulus during resting state (Rest), to an artificial flashing stimulus (Flash), to educational videos which were relatively more engaging (Arith, Fract, StudT), and clips from conventional cinema which were qualitatively most engaging (Wimpy and DesMe, see Stimuli section in Methods for a more complete description). ISC is a stimulus-driven measure of attention (Ki et al., 2016) and it is therefore expected to be indicative of varying levels of engagement (Cohen et al., 2017). A one-way ANOVA determined that ISC significantly depended on the stimulus (F(7) = 78.26, p = 10^−68^; mean +/- STD ISC values: Wimpy: 0.053 +/- 0.036; DesMe: 0.035 +/- 0.023; Arith: 0.019 +/- 0.013; Fract: 0.026 +/- 0.016; StudT: 0.012 +/- 0.009; Flash: 0.030 +/- 0.019; Rest: 0.001 +/- 0.004), indicating that the stimuli significantly varied in engagement level. As expected, ISC in the Rest condition was not significantly different from zero (t-test, t(45) = 0.52, p = 0.4), confirming the notion that ISC reflects stimulus-induced correlations (Dmochowski et al., 2012). A one-way ANOVA was therefore performed on all stimuli excluding Rest, confirming that ISC strongly varies between stimuli (F(6) = 71.70, p = 10^−68^). Tukey post-hoc pairwise comparisons revealed that ISC was significantly stronger when evoked by the qualitatively more engaging stimuli (Wimpy and DesMe), than it was for educational videos (Arith, Fract, StudT; Tukey post-hoc pair-wise comparisons between each pair of videos, Tukey’s HSD: p < 10^−68^). Among the more engaging videos from conventional cinema, Wimpy, a movie trailer for the feature film “Diary of a Wimpy Kid”, evoked a higher level of neural similarity than DesMe, a scene from the animated film “Despicable Me” (Tukey’s HSD: p = 10^−68^). Among the relatively less-engaging educational videos, Fract elicited the highest level of ISC, which was significantly higher than StudT (Tukey’s HSD: p = 10^−68^), but not Arith (Tukey’s HSD: p = 0.2). Interestingly, Arith elicited a level of ISC similar to Flash (Tukey’s HSD: p = 0.5), and the level of ISC elicited by Flash was significantly higher than StudT (Tukey’s HSD: p=10^−68^).

### Intersubject correlation decreases with age

We hypothesized that neural similarity changes with age and therefore examined the correlation between ISC and age. Here, ISC is computed in individuals by measuring the extent to which each subject correlated with the other people in the same stimulus condition. For all of the stimuli excluding Rest, there was a negative relationship between age and ISC (all r’s=-0.68 +/- 0.09, all p’s<10^−68^, Figure 3). ISC did not vary with age during Rest (r = −0.10, p = 0.5, N = 46). This was expected since Rest contained no stimulus to drive EEG signal similarly across subjects.

**Figure 3:**
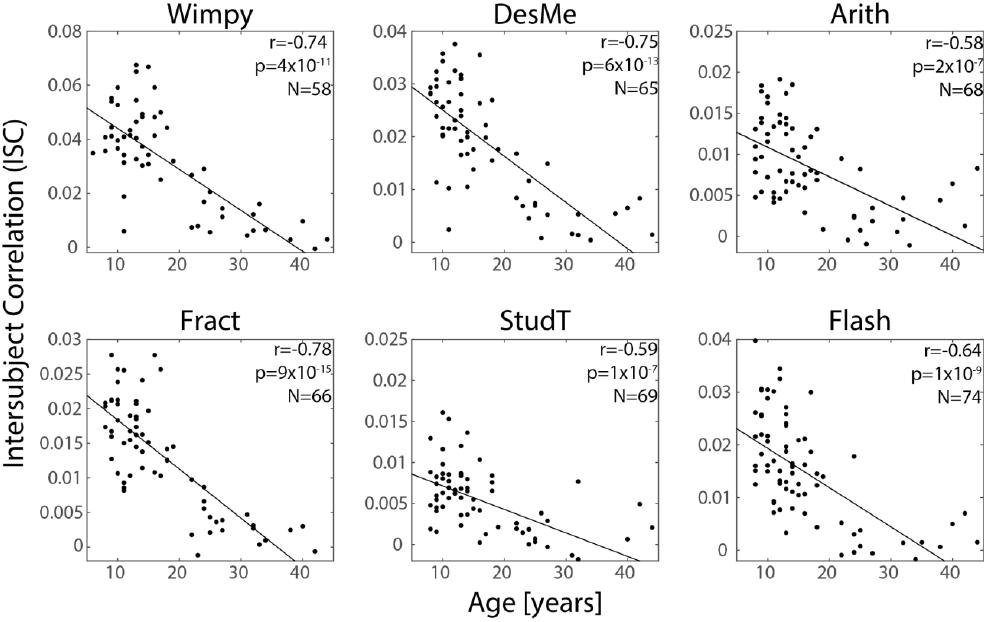
Neural similarity, measured as the intersubject correlation (ISC) of neural activity, decreased with age. ISC was computed for each individual by correlating neural responses from individual subjects with the neural responses from all other subjects for that stimulus (regardless of age and sex). Correlation values ranged from r=-0.58 to r=-0.78, indicating a consistent relationship between maturity and neural variability.

These results indicate that ISC decreases with age. However, most of the subjects in the main study were from the lower half of the age distribution (see Figure 1A). Since the components used to measure ISC are optimized to capture the correlation across all subjects, the components may have been biased by these younger subjects who constituted a majority of the sample. The cohort was therefore divided into two age groups of equal size to eliminate this potential measurement bias. The median split resulted in groups whose ages ranged from 6-14 (mean age 10.74 +/- 2.03) and 15-44 (mean age 23.65 +/- 8.04). The ISC was then recomputed from components extracted separately in each group. A two-way ANOVA with factors of age and stimulus revealed that ISC was significantly modulated by both stimulus (all excluding Rest, F(5, 393) = 63.64, p = 10^−68^) and age (F(1, 393) = 335.46, p = 10^−68^, Figure 4A). For all stimuli, ISC was much higher in the younger age group. To determine whether ISC also decreases within each age group, the correlation between age and ISC was examined separately in each group. In general, ISC decreased with age for several stimuli in both the young group (DesMe: r = −0.35, p=0.03; Arith: r = −0.44, p = 0.006; Flash: r = −0.45, p = 0.02) and the old group (Wimpy: r = −0.45, p = 0.01; Fract: r = −0.41, p = 0.03; Flash: r = −0.71, p = 10^−68^).

**Figure 4:**
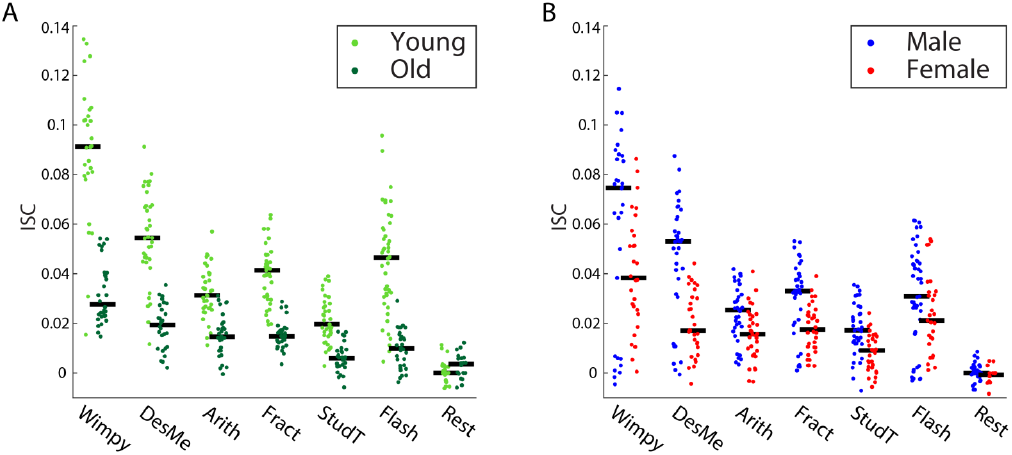
ISC, a measure of neural similarity, was consistently higher among younger ages and males. A: Across all stimuli, ISC was higher for younger subjects (6-14 years, light green) than older subjects (15-44 years, dark green). B: Across all stimuli, ISC was higher for males (blue) than females (red). For both A and B, ISC was computed separately within each age and sex group. Black lines indicate the median.

### Intersubject correlation is elevated in males

Sex is an important factor that influences the developmental trajectory of the human brain (Giedd et al., 1999; Lenroot and Giedd, 2006; Lenroot et al., 2007; Marsh et al., 2008). We therefore explored the relationship between sex and ISC. A two-way ANOVA with factors of sex and stimulus (excluding Rest) revealed main effects for both sex (F(1, 393)=53.11, p = 10^−68^) and stimulus (F(5, 393) = 30.12, p = 10^−68^, Figure 4B). Tukey’s post hoc tests revealed that ISC was consistently higher in males for all stimuli except for Flash where it was marginally significant (Flash: p = 0.06; Wimpy: p = 0.03; DesMe: p = 10^−68^; Arith: p = 0.003; Fract: p = 10^−68^; StudT: p = 10^−68^). To examine whether the sex difference depended on age, the data was separated into four groups with the same age division between 14 and 15 years as above (young-male, young-female, old-male, old-female). ISC was measured within each group and averaged across all stimuli available for each subject to ensure sufficiently large sample sizes (excluding control conditions - Flash and Rest, Figure 5). A two-way ANOVA with sex and age as factors confirmed the age effect (F(1, 87) =98.85, p = 10^−68^), and the sex effect was marginally significant (F(1, 87) = 3.83, p = 0.05). A direct comparison between the sexes in each age group revealed that the sex effect was marginally significant among the young ages (t(53)=2.02, p=0.05, 6-14 years), but not present for the old ages (t(33)=0.28, p=0.8, 15-44 years).

**Figure 5.**
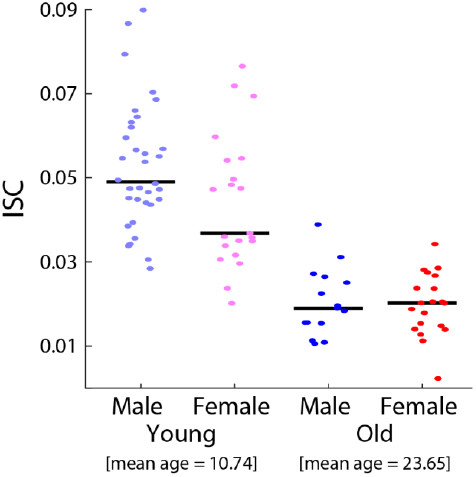
Sex differences in the young disappear with age. Young males were more neurally similar to each other than young females. This sex difference is absent in the older group. Here, ISC was computed within each sex and age group separately and averaged across all stimuli except for Flash and Rest. Black lines indicate the median.

### Age’s effect on inter-subject correlation not due to evoked response difference

The relationships between ISC, age, and sex may be partially driven by the reduction of evoked response magnitude with age (Goodin et al., 1978; Tomé et al., 2015). Although correlation, which ISC measures, is theoretically independent of magnitude, it is possible that a decrease in magnitude corresponds with a decrease in the signal-to-noise ratio, which would result in a smaller ISC. The magnitude of evoked responses was therefore assessed with the Flash stimulus which elicited steady-state visually evoked potentials (SSVEPs, see Methods). SSVEP magnitude weakly declines with age (r=-0.22, p=0.02, N=109, Figure 6A) and two-way ANOVA with age and sex as factors (the same age/sex groups as Figure 5) found the age effect to be marginally significant (F(1,106)=4.00, p=0.05, Figure 6B). There was no significant relationship between sex and SSVEP strength (F(1,106)=3.3, p=0.08)

**Figure 6:**
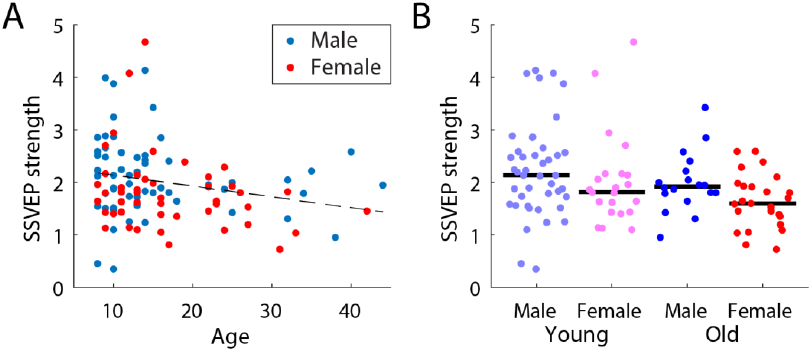
Steady state visual evoked potential (SSVEP) magnitude depended on age, but not on sex. A: SSVEP strength was weakly correlated with age across subjects, but it was no different between males and females. B: SSVEP strength was no different between males and females. Black lines indicate the median.

Since both SSVEP amplitude and ISC decrease with age, we reasoned that SSVEPs could be used to factor out the effect of evoked response strength. Indeed, ISC and SSVEP amplitude are correlated across subjects (r = 0.41, p = 0.0001, N = 84, Figure 7A). To control for the effect of evoked response strength, each individual’s SSVEP amplitude was linearly regressed against ISC, and the portion that could be explained by the SSVEP was subtracted (ISC calculated within the same age/sex group as Figure 5). A two-way ANOVA with age and sex as factors revealed that this residual ISC still significantly varies with age (F(1,81)=85.49, p = 10^−14^), but does not vary with sex (F(1,81)=0.08, p = 0.8, Figure 7B). Additionally, the sex effect is no longer present in the younger group when SSVEP strength is controlled for (t(49)=0.11, p=0.9). The lack of a sex effect may mean that the relationship between sex and neural variability is due in part to evoked response magnitude, but the lack of an effect may also result from the reduced number of subjects for which SSVEP magnitude was available: 84 vs 114. Regardless, neural variability, as assessed by ISC, does increase with age, regardless of the strength of evoked responses.

**Figure 7:**
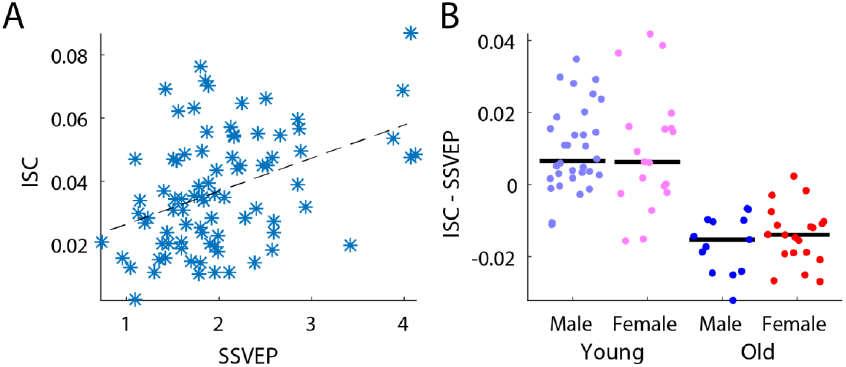
Relationship between ISC magnitude and SSVEP strength. A: SSVEP strength, a measure of the magnitude of evoked responses, correlated with ISC strength, calculated using all stimuli except for Flash and Rest. B: Comparison of ISC strength after SSVEP magnitude was regressed out (ISC - SSVEP) between males and females in the young age and old age groups. While there was a significant difference between the age groups, a difference between the sex groups was not present. Black line indicates the median.

**Figure 8:**
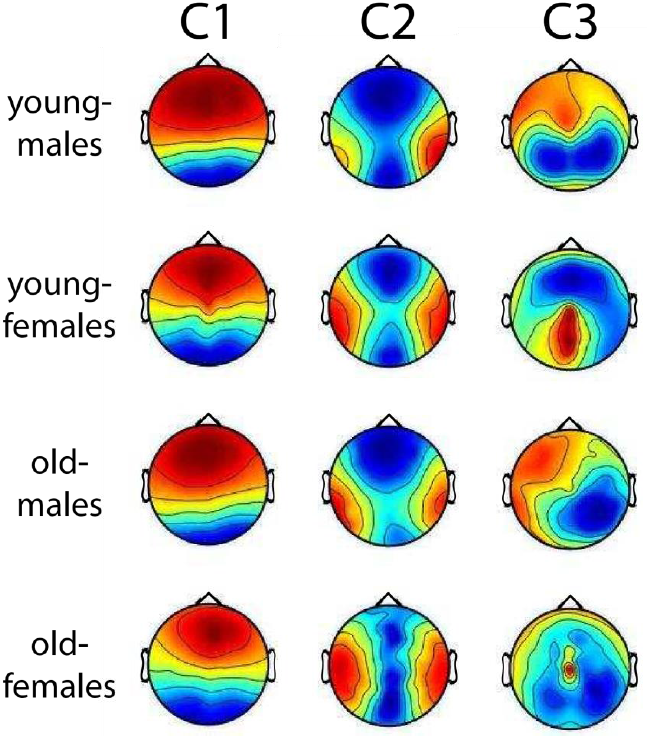
Spatial distributions corresponding to the three strongest components of intersubject correlation (ISC: C1 - C3). Red and blue colors indicate positive and negative correlation of the voltages on the scalp surface with the component activity. These maps are unitless due to an arbitrary scale on the projection vectors. Here, the projections have been computed separately for the combination of the two sex and age groups. As the scalp topographies were relatively consistent across the groups, the differences in ISC across these groups was not due to differences in the spatial topography of correlation within the group.

### Correlated component topographies similar across age and sex groups

ISC was measured using components of the EEG that maximize correlations between subjects. These components are linear combinations of electrodes and can be thought of as virtual sensors (See Methods). To determine if the spatial distribution of the corresponding activity differed across groups, the “forward model” was computed for the largest three components which were used to compute ISC. These component topographies were very similar across all age/sex groups for the strongest two components - C1 and C2 (Figure 7). The third component (C3) was less similar across the groups, but it also constituted a much weaker portion of the ISC (C1 = 0.016 +/-0.009, C2 =0.008 +/-0.005, and C3 =0.004 +/- 0.003, computed as in Figure 3 and averaged across all subjects and stimuli). Thus, for the most part, differences in ISC between age and sex groups were not due to differences in the spatial distribution of neural activity across these groups.

### Replication of results

To confirm these findings, the results were replicated in an independent cohort (N=303) with a reduced stimulus set: Wimpy, Fract, and DesMe. Replicating the results above, ISC also decreased with age in this cohort (Wimpy: r = −0.44, p = 10^−68^, N = 276; Fract: r = −0.37, p = 10^−68^, N = 270; DesMe: r = −0.41, p = 10^−68^, n=281, Figure 9A). A two-way ANOVA with age and condition as factors revealed that ISC is modulated by age (F(1,799) = 35.33, p = 10^−68^) and stimulus (F(2,799) = 272.903, p = 10^−68^, Figure 9B). A two-way ANOVA with sex and stimulus as factors revealed that ISC was also significantly modulated by sex (F(1,823) = 11.12, p = 0.0009, Figure 9C), and stimulus (F(2,823) = 430.95, p = 10^−68^). Finally, a two-way ANOVA which divided the data across age and sex groups replicated the main effect of age (F(1,291) = 17.68, p = 10^−68^), and did not find an effect of sex (F(1,291) = 2.59, p = 0.1, Figure 9D). Follow up analyses that examined a potential sex difference in ISC in each age group revealed that the difference in ISC was present among the young ages (t(224)=2.29, p=0.02, 5-14 years), but not the old ages (t(67)=0.59, p=0.6, 15-21 years). When the median was calculated according to the median of the replication distribution (split at 10/11 years, see Figure 1B for age distribution), the above results were unchanged. In summary, all results from the main experiment replicated in this independent cohort.

**Figure 9:**
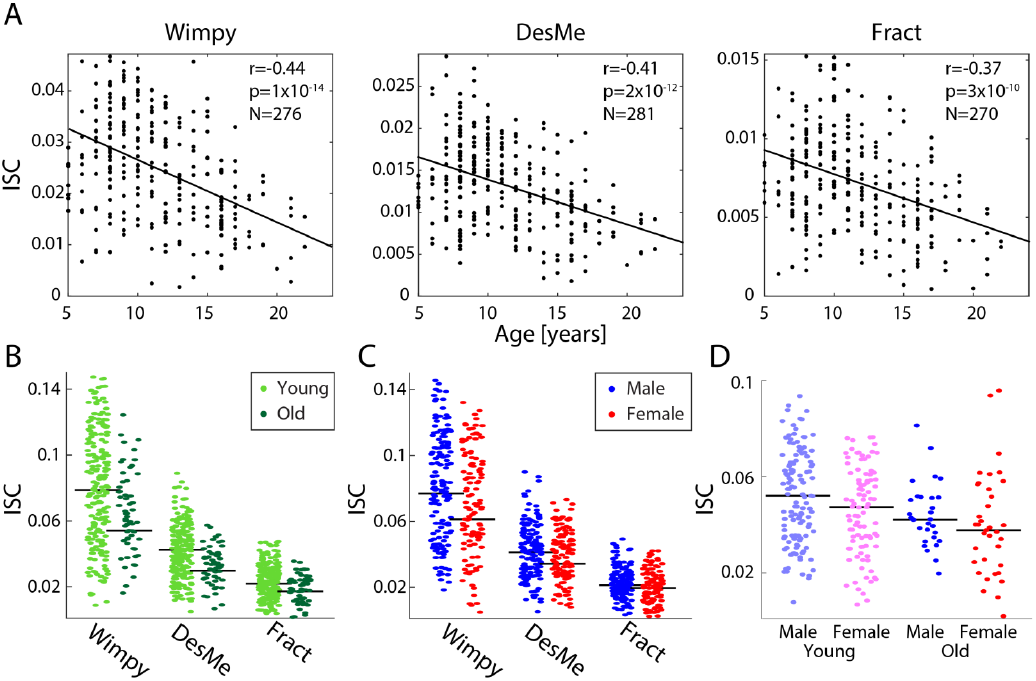
The results from the main study replicated in an independent cohort (N=303). A: ISC decreased with age in the replication cohort. ISC was computed for each individual by correlating responses from individual subjects to those from all other subjects (regardless of age and sex) for that stimulus. Correlation values ranged from r=-0.37 to r=-0.44. Note that for every stimulus a different number of subjects was available. B: Across all stimuli, ISC was higher for younger subjects (6-14 years, light green) than it was for older subjects (15-44 years, dark green) in the replication cohort. For consistency, the split between the ages was consistent between this study and the main study C: Across all stimuli, ISC was higher for males (blue) than females (red) in the replication cohort. For both B and C, ISC was computed separately within each age and sex group. Black lines indicate the median. D: Sex differences in the young disappeared with age in the replication cohort. Young males were more neurally similar to each other than young females, and this sex difference was absent in the older group. Here, ISC was computed within each sex and age group separately and averaged across all stimuli used in the replication cohort. Black lines indicate the median.

## Discussion

The present work demonstrated age- and sex- related variability among individuals with respect to their neural responses to complex naturalistic stimuli. Specifically, ISC was significantly correlated with age for both naturalistic videos and artificial visual flashes. Younger subjects (6-14 years) exhibited less variable neural responses than older subjects (15-44 years). A parallel finding revealed that young males exhibited more similar responses to the stimuli than young females, a difference which was only present in the younger cohort. These age and sex effects may result from neural development, consistent with the notion that neural maturation occurs later in males than in females (Lenroot et al., 2007; Marsh et al., 2008; Mous et al., 2017). A qualitative analysis of the spatial distribution of the correlated activity revealed that the observed age and sex differences are largely driven by the same neural components, lending more weight to the idea that the observed differences in age and sex stem from a common developmental feature. Finally, a replication study with 303 participants yielded similar results.

A possible confound for the present results is that the neural correlations found across subjects are due to correlations in overt behaviors such as eye movements. However, it is unlikely that eye movements follow the same developmental trajectory as neural responses because eye movement trajectories evoked by videos actually become more similar with age (Kirkorian et al., 2012). Thus, although the gaze patterns evoked by videos seem to converge with maturity, potentially driving similar bottom-up neural processes, neural similarity as measured by ISC, *decreases* with age. This indicates that patterns of neural activity may potentially increase in their diversity with age as top-down factors relating to the interpretation of naturalistic stimuli develop. Even in the condition where subjects were instructed to maintain a fixed gaze position (Flash), ISC decreased with age. Future studies with fine-grained eye-tracking during EEG could more definitively answer this question.

The observed ISC magnitude changes with age and sex may also be partially dependent on evoked response magnitudes which typically decrease with age. While the amplitudes of auditory event related potentials and their components decline with age (Goodin et al., 1978; Tomé et al., 2015), other components increase with age (Dinteren et al., 2014), or remain stable across development (Kujawa et al., 2013). Although correlation, as measured by ISC, is in principle insensitive to magnitude, it is possible that weaker stimulus evoked responses in adults may be overpowered by non-stimulus related neural activity (i.e., “noise”) (Hammerer et al., 2013). In this case, a smaller fraction of the signal would correlate across adults in comparison to children. To control for the effect of age, the magnitude of steady state visual evoked potentials (SSVEPs) was regressed from the ISC. The result indicates that SSVEP amplitude cannot explain the age effect, but it may explain the sex effect, indicating that males have stronger evoked responses than females (Figure 4B and 6B). However, it is worth noting that ISC and SSVEPs measure very different facets of neural activity. SSVEPs, extracted from early visual processing areas in V1, likely represent low-level visual processes. ISC, on the other hand, may be driven by higher level cortical areas since the spatial distributions of the two dominant components (Figure 8) do not resemble low-level sensory evoked responses. Parallel work indicates that the first component (C1), which captures the majority of the correlated activity, is a supramodal component that is driven by both auditory and visual stimuli (Cohen and Parra, 2016).

It is also possible that ISC decreases with age because adults process the world with more diverse brain activity. In this view, adults have more highly variable stimulus-evoked responses and their neural activity is therefore less similar across subjects. In this case, it would be likely that the dimensionality of neural responses, a measure of their complexity, increases with age (Anokhin et al., 2000; Mcintosh et al., 2008; Vakorin et al., 2013). A clear trend, indicative of a difference in the dimensionality between either the young and old group, or males and females, was not apparent in these data.

The literature related to developmental changes in oscillatory power is mixed. While most studies in children report a decrease in power with age for most frequency bands (Gasser et al., 1988; Harmony et al., 1990; Clarke et al., 2001), some studies report an increase in power during a task (Liu et al., 2014), or in selective frequency bands, e.g. alpha (Benninger et al., 1984) or gamma (Uhlhaas et al., 2009). Our data suggest that these differences may relate to whether activity was recorded during a stimulus or task. It may be that the mature brain requires fewer resources to process the world to which it has grown accustomed. This finding may underlie the weaker and more divergent neural patterns found in the ISC of older brains (Campbell et al., 2015).

The idea that maturity is marked by variability is not new (Campbell et al., 2015). It aligns with theories from neural systems modeling and human studies (Mcintosh et al., 2008; Vakorin et al., 2011). In these models, moderate amounts of noise or variability facilitate efficient responses in complex environments. Increased variability may be the reason for reduced evoked response magnitudes since event related potentials are obtained by averaging across many events that are inherently sensitive to signal noise. It is therefore possible that the increased variability of evoked responses across trials with age results in reduced ERP magnitudes.

In the age range examined, neural development is a dynamic process. At the macro level, longitudinal structural neuroimaging shows that cortical thinning occurs from childhood through early adulthood, progressing in a caudal to rostral pattern (Gogtay et al., 2004; Giedd et al., 2015). At the micro level, synaptic pruning and myelination, particularly in the frontal lobe, are ongoing during this period (Rakic et al., 1994; Huttenlocher and Dabholkar, 1997; Cox et al., 2016). From a functional perspective, studies of functional connectivity and task-based fMRI suggest that functional maturation tends to follow a “diffuse to focal pattern” (Durston et al., 2006; Grill-Spector et al., 2008; Fair et al., 2009; Kelly et al., 2009), and may correspond to the extraordinary advances in behavior during childhood (Xiao et al., 2016). Speculatively, the decreased ISC strength in older ages may reflect greater inter-individual variability that results from the interplay of structural and functional “streamlining” of neural architecture with distinct life experiences (e.g. cortical thinning, synaptic pruning and diffuse-to-focal shifts in functional patterns). However, one limitation of the present study is that it is cross-sectional rather than longitudinal, it is therefore difficult to make developmental claims based on the age-based differences demonstrated here (Kraemer et al., 2000).

The age-related effect, may also be echoed by the sex difference in neural variability. Longitudinal studies have demonstrated that females mature prior to males in a range of anatomical measures (Lenroot et al., 2007; Lim et al., 2015). However, differences in developmental trajectories between males and females may be complicated by the fact that the sexes ultimately differ in their mature neuroanatomy (Marsh et al., 2008). Here, sex-related differences in neural variability were only seen among younger subjects, suggesting that this is a development-related difference. Prepubescent and early teenage years are marked by sex differences in behavioral maturity that may not be present in later years (Mous et al., 2017). The difference in neural variability may also be due to pubertal stage since it is known that females reach pubertal maturity 2-3 years prior to males (Sisk and Foster, 2004). However, physiological pubertal stage was not measured here, and it is therefore not possible to determine whether the sex differences were related to this factor.

Among the different stimuli used, the clips from conventional cinema (Wimpy and DesMe) evoked a higher level of ISC than the educational videos (Arith, Fract, and StudT). The Hollywood clips were rich with scene cuts and dynamic visual cues and are therefore expected to elicit strong levels of ISC (Poulsen et al., 2017). However, previous research has also shown that engagement with narrative stimuli modulates ISC, and it is therefore likely that these Hollywood clips are more effective at engaging attention and thus elicit stronger ISC (Dmochowski et al., 2014; Ki et al., 2016; Cohen et al., 2017). Although the ISC differences between age and sex may be due to each cohort’s average level of attention, no independent measures of engagement or attention were collected. It is therefore not possible to determine whether the present effects are driven by attention or differences in low-level stimulus features. However, it is true that most of the videos were aimed at younger audiences (i.e., *Despicable Me, Diary of a Wimpy Kid*) and older subjects may have therefore been less interested. Future work may benefit from looking at objective measures of engagement (Cohen et al., 2017) in the different cohorts studies here. An understanding of such factors, and their impact on behavior, may be of relevance to models of media-based addiction (e.g., internet addiction, pornography addiction), as well as commercial neuroscience enterprises. Regardless, it is of note that the age effect seen for the naturalistic videos was echoed in the SSVEP condition. Since this stimulus should be equally (un)engaging for all ages, this favors an interpretation based on neural maturation rather than attention.

Future work should recruit a larger sample of subjects above age 15 to determine whether the age-related decline in ISC observed in later teenage years continues in adulthood, or might even reverse in older age (Grady, 2012; Campbell et al., 2015). Future studies with clinical cohorts could explore the potential link between ISC and behavioral markers of neural development. It is possible that neural variability is not only a marker of maturity, but it is also a marker of neuropsychiatric disorders (Dinstein et al., 2015). The methods used here provide a novel way of assessing such markers under complex, naturalistic conditions.

Overall, the current results regarding intersubject correlation in children and adults are interpreted in the context of neural maturation. Although males are delayed in the development of the neural variability that appears to be a mark of maturity, the data presented here indicate that with normal development they are no different than females as adults. Thus, with maturity, neural function becomes more variable.

## Acknowledgements

We thank Stefan Haufe for suggesting code to compute ISC using symmetrized between- and within-subject covariances.

## Author contributions

MM, LP, SC and NL designed the study. AP, SC, LP, and LA analyzed the data. All authors contributed to the manuscript writing.

